# Phytochemical and Therapeutic Evaluation of *Ficus nervosa* Leaves: Impact on Oxidative Stress, Analgesic, Diarrhea, and Blood Clotting in a Mice Model

**DOI:** 10.1101/2024.11.19.624432

**Authors:** Zubair Khalid Labu, Samira Karim, Rahima Akter, Kaniz Fatema, Sarder Arifuzzaman, Tarekur Rahman, Md. Imran Hossain, Farhina Rahman Laboni

## Abstract

**Background:** *Ficus nervosa* (FN) leaves have been traditionally used in ethnomedicine for the treatment of various ailments, this study aimed to investigate the cold methanol extract of FN for its antioxidant, analgesic, antidiarrheal, and thrombolytic properties, along with preliminary phytochemical screening to identify key bioactive compounds and acute toxicity test to assess the safety profile of the extract.

**Methods:** In this study, we conducted an initial investigation to identify the major phytochemical groups present in the leaves of *Ficus nervosa*. Using conventional phytochemical screening methods on cold methanol extracts, we consistently identified phenols and flavonoids as the predominant bioactive compounds.

Following this phytochemical characterization, we assessed the biological activities of the extracts across multiple models. The antioxidant capacity and free radical-scavenging ability of the leaves extracts were evaluated using the DPPH (2,2-Diphenyl-1-picrylhydrazyl) assay. Analgesic efficacy was assessed through acetic acid-induced writhing and electrically induced heat nociception tests. Antidiarrheal effects were examined using castor oil-induced diarrhea and an enteropooling assay in mice. To determine thrombolytic activity, we conducted a human blood clot lysis assay, while preclinical toxicity testing was conducted to evaluate the safety profile of the extracts.

Statistical analysis was performed using one-way ANOVA with Duncan’s Multiple Range Test (DMRT) to validate the significance of the results.

**Results:** The study of Antioxidant Potential, the ESF’s (ethyl soluble fraction) antioxidant capacity, measured at 409.91 ± 0.59 mg of ascorbic acid equivalent (AAE)/g in the phosphomolybdenum assay, signifies a strong free radical scavenging activity. This level of antioxidant action, alongside a potent free radical neutralizing concentration (IC₅₀ of 9.99 ± 0.41 μg/mL), suggests that the ESF could effectively counteract oxidative stress. Analgesic activity at 500 mg/kg, FN extract produced a 68.05% suppression of writhing in mice, which was statistically significant (p<0.001) compared to standard morphine’s (74.05%) inhibition. The extract also significantly prolonged reaction latency to thermal-induced pain in hotplate model. The antidiarrheal activity showed 52.09% and 59.0% suppression of defecation, compared to standard loperamide’s 73.1%, with statistically significant p-values (p<0.04 and p<0.01). These effects, combined with the notable decrease in intestinal fluid accumulation and defecation frequency in enteropooling assays, indicate that FN extract might interact with intestinal motility or fluid secretion pathways, providing antidiarrheal benefits. Additionally, the chloroform-soluble fraction of FN exhibited moderate thrombolytic activity, achieving 39.79% clot lysis compared to streptokinase (69.52%). Although this thrombolytic potential is moderate, it indicates that the fraction contains compounds with some fibrinolytic properties, which could be useful as adjunctive agents in preventing thrombosis.

**Conclusion:** Overall, the cold methanol extract of FN leaves demonstrates the therapeutic potential in preclinical settings. Future research is warranted to isolate the specific bioactive compounds and explain their mechanisms of action to further support the development of new treatments and contributing to modern medicinal practices based on this plant leaves.

## 1. Introduction

Plant-based natural compounds are utilized globally as complementary and alternative medicine, improving people’s health and well-being. Lots of medicines we use, like aspirin, digoxin, morphine, ephedrine, quinine, tubocurarine, and reserpine, originally came from medicinal plants. Phenols and flavonoids are desirable phytoconstituents due to their hydroxyl groups, which can decompose peroxides, repair oxidative damage, and quench singlet and triplet oxygen [1]. Reactive oxygen species (ROS) and free radicals are released by cells and have the power to destroy individual cells causes heart disease, alzheimer’s, parkinson’s, cancer, and diabetes are among these conditions [2]. Malignant cells release a range of reactive oxygen species (ROS) that aid in the progression of cancer as a result of the oxidation of RNA, protein, DNA, and other macromolecules [3]. It has been shown that phenolic and flavonoids can inhibit or delay cancer by reducing oxidative damage and promoting RNA and DNA repair [4]. Heart attacks, stroke, ischemia, clot formation, and other potentially lethal conditions are brought clot formation in the cardiovascular system [5]. Clot formation has been the leading cause of death in recent times. Among the clot break down drugs currently in use are plasminogen activator, streptokinase, and urokinase; nevertheless, all of them have negative side effects [6]. Novel or non-traditional drugs are required for their care, there is a positive clot break down effect associated with plants high in phenols [7]. Research has indicated that the anti-diarrheal properties of flavonoids and polyphenols are clearly present. Anti-diarrheal effect may be attributed to tannins, saponins, and terpenoids [8]. Leaves extract significantly decreased intestinal fluid flow and defecation in the castor oil and normal saline enteropooling test and also castor oil-induced diarrhea test, respectively, it is believable to conclude that the extract has antidiarrheal effect.

*Ficus nervosa* (Moraceae) is a monoecious, evergreen, medium-sized tree that can grow up to 35 meters tall. It is commonly found across various parts of Asia, particularly at low elevations in Taiwan, Malaysia, India, Bangladesh, and South China, and has been traditionally used to treat diabetes, hemorrhagic conditions, infections, and rheumatism [9]. The leaves, measuring 0.08–0.2 m in length, are evergreen, glabrous on both sides, have a coriaceous texture, and feature a complete border. The plant is known for its traditional applications in treating diabetes, antihemorrhagic conditions, rheumatism, infections, and ulcer disorders [10].

An ethnopharmacological evaluation of *F. nervosa* is warranted due to its established traditional medicinal uses, potential for the discovery of bioactive compounds, the opportunity to address scientific gaps, and the promotion of conservation and sustainable use. This research holds promise for uncovering new therapeutic applications and bridging traditional knowledge with modern medicine. *F. nervosa* is rich in bioactive compounds such as flavonoids, phenolic acids, tannins, terpenoids, alkaloids, coumarins, and lignans. These compounds exhibit a broad spectrum of therapeutic activities, including antioxidant, anti-inflammatory, antimicrobial, and anticancer properties [11]. Literature surveys indicate that *F. nervosa* contains numerous secondary metabolites. Traditionally, its fresh sap has been used as a local hemostatic agent to treat external wounds, with its wound-healing, antidiarrheal, and hemostatic properties widely documented [12].

Tannins, known for their astringent properties, contribute to the ability of *F. nervosa* to precipitate proteins, forming a protective layer that reduces bleeding and prevents infections. Flavonoids, abundant in this species, are polyphenolic compounds with antioxidant, anti-inflammatory, and antimicrobial activities. The hydroxyl groups in flavonoids enhance their ability to neutralize free radicals, repair oxidative damage, and facilitate wound healing [13].

In the Asian subcontinent, aqueous extracts of *F. nervosa* leaves are traditionally employed for treating wounds and alleviating pain [14]. Phytochemical investigations into this plant have drawn significant interest due to its medicinal history and bioactive composition [15,16]. Recent studies have focused on evaluating its phytoconstituents and medicinal potential, particularly its antioxidant, antibacterial, antidiarrheal, and hemostatic activities.

The traditional use of *F. nervosa* for managing oxidative stress, analgesic, diarrhea, and blood-related disorders is well-documented, particularly in Mirjapur village in the Tangail district of Bangladesh, where its sap is used as a local remedy for various ailments [17]. In this study, Pearson correlation analysis was used to explore the relationships between the phenol, tannin, and flavonoid content of *F. nervosa* leaves and their **o**xidative stress, algia, diarrhea and blood clotting. Methanol extracts from the leaves were evaluated, revealing significant amounts of these bioactive compounds.

The purpose of this work is to investigate the phenolic and flavonoid content of *F. nervosa* leaves and assess their relationship, using a Pearson correlation analysis, with antioxidant, antidiarrheal, analgesic, and thrombolytic properties. In order to assess these activities, we made cold methanol extracts from *F. nervosa* leaves together with their five organic solvent soluble fractions and evaluated such activities, and interrelated them with the phenol and flavonoid contents.

## 2. Materials and Methods

### 2.1 Sampling and proper documentation

Plant leaves were collected from Mirjapur village in the Tangail district of Bangladesh (geographical coordinates: 24.05° N, 89.92 ° E) is where the plant leaves were collected. After completing the identification of plant leaves by Bangladesh National Herbarium’s taxonomist followed by provided taxonomical accession number # 454089 and with all necessary permissions obtained to continue our study. The experimental research and field studies involving *F. nervosa* (L.) leaves strictly adhered to established international research guidelines, including the Convention on Biological Diversity (CBD) and the study also received approval from the Institutional Review Board (IRB) for Plant Research, ensuring that all research activities were conducted ethically and responsibly. The collection process was carried out sustainably, with careful measures in place to prevent any detrimental impact on the local ecosystem. This approach not only complied with all ethical standards but also supported the preservation of biodiversity and environmental integrity.

### 2.2 Chemicals and Reagents

Analytically graded chemicals were used throughout the study, including methanol (liquid chromatography grade, ≥99.8%), ethyl acetate (≥99.9% GC), pet-ether (≥80%), chloroform (≥99% ACS Reagent Grade), carbon tetrachloride (≥99.9%), gallic acid (98%), catechin (≥99.8%), Streptokinase (SK), lyophilized SK that is sold commercially was utilized. Folin-Ciocalteu reagent (standard reagent grade), and aluminum chloride (anhydrous sublimed, ≥99.8%), all provided by Science Park Chemicals Ltd., Bangladesh.

### 2.3 Equipment and Instrumentation

To ensure precision and minimize experimental errors, all instruments were calibrated according to standard procedures. The equipment used in the study included:

- UV-Vis spectrophotometer (Model: UV-1700 series)
- Casio digital stopwatch (Model: HS-70W-1DF)
- Mechanical dryer (Model: CG-CG23KW-150KW)
- Digital analytical balance (Model: PS. P3.310)
- Thermostat water bath (Model: HHW21.420AII)
- Autoclave (Model: DSX-280KB)
- Rotary evaporator (Model: PGB002)
- Vortex mixer (Model: VM-11)

### 2.4 Drying and grinding of plant leaves

The collected plant leaves were separated from unwanted materials, thoroughly washed with water to remove any residual dirt, and sun-dried for one week. Subsequently, they were further dried in a mechanical dryer. The dried leaves were then mechanically processed into a coarse powder, stored in an airtight container, and kept in a cool, dry, and dark place until analysis.

### 2.5 Plant material extraction

A total of 500 grams of powdered *Ficus nervosa* leaves material was placed in a clean, dark-colored glass container with a flat bottom to minimize light exposure, which is known to reduce glare, improve sample stability, and block UV rays. The powder was submerged in 1.5 liters of methanol at a controlled temperature of 25°C. To prevent air exposure, the container was tightly sealed, and the contents were left to extract over ten days with periodic shaking and stirring to enhance the extraction efficiency.

After the extraction period, the solution was decanted and initially filtered through cotton to remove larger particles. This was followed by filtration through Whatman No. 1 filter paper to ensure purity. The resulting filtrate was concentrated at 40°C using a rotary evaporator, yielding a sticky, greenish-black substance, identified as the crude methanolic extract [18].

Approximately 12 grams of the crude methanolic extract were then dissolved in high-purity (≥99.8%) methanol. The dissolved extract underwent Kupchan partitioning to separate it into distinct fractions based on solubility properties, following standard fractionation techniques [19]. After complete evaporation, the following fractions were obtained: Ethyl acetate-soluble fraction (ESF):4.2 grams, Petroleum ether-soluble fraction (PSF): 1.0-gram, Chloroform-soluble fraction (CSF):5.0 grams, Carbon tetrachloride-soluble fraction (CTSF):1.5 grams, Methanol-soluble fraction (MSF):0.2 grams, Aqueous fraction (AQF):0.1 grams.

Each fraction was subsequently stored at low temperatures to preserve stability for further biological analyses.

### 2.6 Phytochemical screenings

The confirmatory qualitative phytochemical screening of the crude extracts was conducted to detect the primary classes of compounds, including tannins, saponins, flavonoids, alkaloids, phenols, glycosides, steroids, reducing sugars, and carbohydrates, using established standard protocols [20]. To verify the presence of primary phytochemical classes in crude extracts, we conducted a confirmatory qualitative phytochemical screening. This analysis targeted compounds such as tannins, saponins, flavonoids, alkaloids, phenols, glycosides, steroids, reducing sugars, and carbohydrates, following standard protocols. Although previous studies have identified various phytochemicals in *F. nervosa* and related species, re-confirmation in our specific samples is essential. Variations in environmental conditions, soil composition, and geographical factors can affect the concentration and even the presence of certain phytochemicals. Hence, re-screening is not only crucial for validating previous findings but also for potentially detecting minor or previously overlooked compounds that may contribute to the observed biological activities [21]. This thorough screening ensures that any therapeutic or biological effects are accurately attributed to the phytochemicals present in our sample from the particular collection site, laying the groundwork for further quantitative analyses and biological evaluations.

### 2.7 Determination of total phenolic content (TPC)

With a few minor modifications, the recommended methodology was applied in the experiment to determined TPC using the Folin-Ciocalteu reagent [22]. Two milliliters of crude methanolic extract and all of its organic soluble components were liquefied with two milliliters of distilled water to achieve the final concentration of one milligram per milliliter. A mixture of 0.5 mL of extractives (1 mg/mL) and 2 mL of diluted Folin-Ciocalteu (previously diluted 10-fold with deionized water) was placed in the test tube. After allowing the mixture to stand at 22 ± 2 °C for five minutes, 2.5 milliliters of 7.5% Na_2_CO_3_ was added to each combination. The mixture was gently stirred for 20 minutes without allowing any substantial jerking to allow for color development. The UV-Vis spectrophotometer (Model: UV-1700 series) was used to detect the color shift’s intensity at 760 nm. The absorbance value indicated the compound’s TPC. To evaluate the extract and standard samples, a final concentration of 0.1 mg/mL was employed (20). To quantify TPCs as mg of GA/g of dry extract, the GA equivalent, or GAE using the calibration curve.

### 2.8 Determination of total flavonoid content (TFC)

The aluminum chloride (ALCL_3_) colorimetric method was utilized to ascertain the total flavonoid content of the crude methanolic extract and its partitionates, which include PSF, MSF, ESF, CTS, and CSF [23]. To summarize, 0.1 mL of 10% AlCl_3_ and 1.5 mL of crude extract (1 mg/mL methanol) and its partitionates were combined, and 0.1 mL of 1 M Na-acetate was then added to the reaction mixture. The mixture was left to stand for half an hour. Next, 1 mL of a 1 mol/L NaOH solution was added, and the combination was finally brought to a final volume of 5 mL using double-distilled water. Following a 15-minute incubation period, at 415 nm, absorbance was measured. Using a calibration curve, the total flavonoid content was determined and reported as mg of catechin equivalent per g of dry weight. The total flavonoid content was ascertained using the calibration curve.

### 2.9 Antioxidant activity

More than a hundred human health disorders are linked to free radicals, including ailments like arthritis, ischemia, atherosclerosis, and reperfusion injury to several tissues, harm to the central nervous system, cancer, gastritis, and acquired immune deficiency syndrome [24]. The leaves extracts’ antioxidant capacity and capacity to scavenge free radicals on the stable radical DPPH (2,2-Diphenyl-1-picrylhydrazyl) were determined by applying the method described by Williams *et* al. [25]. The method is typified in the presence of an antioxidant that donates hydrogen when the radical DPPH in methanol solution is reduced as a result of the non-radical DPPH-H being produced. The spectrophotometrically amplified color change in methanol from purple to yellow showed that DPPH was effective at scavenging free radicals. Together with 2.0 mL of crude methanolic extract and its extract fractions at different concentrations, 3.0 mL of DPPH methanol solution (20 µg/mL) was added. The extract was found to be able to bleach a purple-colored methanol solution containing DPPH radicals when its antioxidant capacity was compared to ascorbic acid (AA) using a UV spectrophotometer. AA was subjected to positive control. After calculating the amount of AA to dissolve in methanol, a mother solution containing 1000µg/mL was produced. Differential concentrations ranging from 500.0 to 0.977 µg/mL were obtained by progressively diluting the mother solution. The determined quantity of different extracts was dissolved in methanol to create the mother solution, which contained 1000 µg/mL. The mother solution was gradually diluted in order to achieve differential concentration. After weighing and dissolving 20 mg of DPPH powder in methanol, 20 µg/mL of DPPH solution was produced. The amber reagent bottle and the prepared solution were kept in the light-proof box. The sample (extractives/control) methanol solution was mixed with 2.0 mL of DPPH methanol solution (20 µg/mL) at different concentrations (500 µg/mL to 0.977 µg/mL). After a 30-minute reaction at room temperature in a dark atmosphere, the absorbance was measured at 517 nm using a UV spectrophotometer using. To calculate the percent (I%) of free radical DPPH inhibition, the following formula was used:

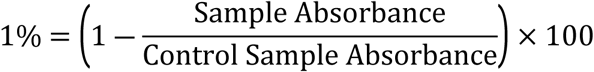

To determine the % inhibition, a similar process was carried out using standard AA rather than the sample solution (leaves extract). The extract concentration that would produce 50% inhibition (IC_50_) was found using the graph that plots the proportion of inhibition against extract concentration.

### 2.10 Ethical Clearance

This study was carried out in strict accordance with the recommendations in the Guide for the Care and Use International Centre for Diarrheal Disease Research, Bangladesh (ICDDRB) guidelines, which was followed for conducting animal experiments. The World University of Bangladesh Ethical Committee Ethics of Animal Experiments read out the study through and gave its approval (Approval no # WUB/2023/249L11) to continue our study. The Ethical Committee on Animal Experiments at the World University of Bangladesh, in collaboration with our research team, mandated that the experimental mice used in this study would not be reused for any subsequent experiments. Upon completion of the study, all were promptly and humanely euthanized to minimize distress, fully in accordance with ethical guidelines. We strictly adhered to humane euthanasia methods and environmentally responsible handling practices, ensuring that we met the highest standards of animal welfare and ethical responsibility.

### 2.11 Experimental animal care

For this experiment, 8–9-week-old Swiss albino mice, weighing 80–90gram on average were used. Mice were housed under controlled environmental conditions, with a temperature maintained between 20°C and 24°C, relative humidity at 30%, and a photoperiod of 12 hours of light followed by 12 hours of darkness. The animal rooms and cages were regularly cleaned, and suitable nesting material was provided within the cages to enable the mice to regulate their microclimate effectively, in addition allow them to perform their natural nesting behaviors. They were given Jahangir Nagor University; Bangladesh formulated food ad-libitum and tap water. Mice kept in a typical setting for a week at the World University of Bangladesh research laboratory to help them acclimate.

### 2.12 Method of sacrifice

The method of cervical dislocation for euthanizing mice under 200 grams was performed without anesthesia, adhering to the 2020 AVMA and IACUC guidelines [26]. Trained personnel carried out the procedure skillfully, using just the right amount of force to ensure the dislocation happened quickly and effectively, minimizing any pain or distress. During cervical dislocation, the thumb and index finger were positioned on either side of the animal’s neck at the base of the skull while the animal lay on a table surface. On the other hand, the base of the tail is firmly and steadily pulled to cause separation of the cervical vertebrae and spinal cord from the skull. Following the procedure, we monitored the mice for over 60 seconds until the heartbeat ceased before disposal. Trained personnel monitored the mice for signs of pain or distress throughout the process. Critical assessment of the presence or absence of pain and/or distress between control and drug treated mice behavior has been taken carefully. To prevent any environmental contamination, the carcasses were wrapped in a plastic bag, placed in a box, and buried underground. When choosing a burial site, we made sure there were no nearby underground water sources or flood-prone areas to prevent water contamination. We also looked for soil that wasn’t too sandy [27].

### 2.13 Acute toxicity test

For this experiment, 8–9-week-old Swiss albino mice, weighing 80–90gram on average were used. The acute toxicity test was carried out in accordance with OECD (Organization for Economic Co-operation and Development) Test no. 423: acute oral toxicity-acute toxic class method, 2002 with minor modifications was employed to evaluate the acute oral toxicity of *F. nervosa* methanol leaves extracts [28]. The extract was administered carefully through feeding needle to mice by gavage or intraperitoneal route at different concentrations of 200, 400, 800, 1600, and 3200 mg/kg in order to evaluate the acute toxicity test [29]. Prior to administering extract to each mouse of groups (six mice in each group), were kept for fasting condition till 16 hours. Following treatment, the animals were continuously monitored for 1 hour, then intermittently for 4 hours, and subsequently observed over a 24-hour period to assess any behavioral changes, signs of toxicity, or symptoms of death. The latency to death was tracked for 14 days. All mice had unrestricted access to food and water throughout the study, and acute toxicity was evaluated by systematically recording any changes in daily food intake, water consumption, body weight, mortality rates, and visual observations over a 48-hour monitoring period [30]. The LD_50_ value was determined by calculating the geometric mean of the lowest dose that caused death and the highest dose for which the animal survived [31].

### 2.14 Analgesic activity test

In the current investigation two different methods were used for testing the possible peripheral and central analgesic effects of *Ficus nervosa* leaves; namely acetic acid induced writhing test and hot plate test in mice respectively.

#### 2.14.1 Acetic acid induced writhing in mice

The analgesic activity of the crude methanolic extract of FN leaves was investigated through the acetic acid-induced writhing test in mice, a well-established method for assessing peripheral analgesic effects as initially outlined by Koster *et* al. [32]. The animals were divided into four groups including control (Group I), positive control (Group II) and two test groups (Group III-IV). The animals of group III and IV were administered test substance at the dose of 250 and 500 mg/kg body weight respectively. Positive control group received diclofenac (standard drug) at the dose of 25 mg/kg body weight and negative control group was treated with 1% Tween 80 in distilled water at the dose of 10 ml/kg body weight. Test samples, standard drugs and negative control vehicle were administered orally 30 min before intraperitoneal administration of 0.7% acetic acid. After 15 min of time interval, the writhing (constriction of abdomen, turning of trunk and extension of hind legs) was observed on mice for 5 min. Acetic acid activates the pain nerve and is used to simulate writhing by releasing endogenous chemicals [33]. Each mouse’s writhing number (squirms) was accurately counted for fifteen minutes.

The percentage inhibition of writhes was calculated using the following formula:

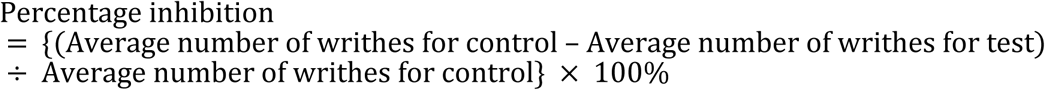

#### 2.14.2 Hot plate test in mice

"The extract of FN leaves was screened for centrally acting analgesic activity using the method outlined by Eddy and Leimbach [34]. The temperature of the hot plate was maintained at 45±0.5⁰C. Only animals which responded when placed on the hot plate within a period of 30 s were selected for the experiment. Thirty of the selected mice were randomly divided into five groups consisting of six mice each. Groups I and II were orally treated with distilled water (10 ml/kg negative control) and morphine (5 mg/kg positive control) respectively. Groups III and IV were orally pre-treated with crude methanolic extract doses of 250, 500 mg/kg respectively. A cut off time of 20 s was used to avoid paw damage. One-hour post treatment, the time taken for each animal to either jump, lick or flutter their paws was taken as the reaction time and recorded at 0, 30, 60, 90 and 120 minutes.

### 2.15 Anti-diarrheal activity

In this study, the anti-diarrheal activity of *Ficus nervosa* leaves was evaluated using two methods: castor oil-induced diarrhea in mice and the enteropooling assay. The two methods provide insights into different mechanisms of action. Castor oil-induced diarrhea in mice typically evaluates the ability of a compound to reduce intestinal motility and secretion, while the enteropooling assay examines the inhibition of fluid accumulation in the intestines. Together, these methods help clarify whether the leaves act by reducing bowel movement, fluid secretion, or both.

#### 2.15.1 Castor oil induced diarrhea in mice

Mice capable of inducing diarrheal activity were identified through an initial screening process before the test commenced, the procedure was conducted in accordance with the description provided by Karim *et* al. [35]. The groups of mice, each consisting of six mice were formed. Group under control, given a vehicle (1% Tween 80 in water) orally at a dose of 10 mL/kg body weight; the positive control group received an oral dose of 3 mg/kg body weight of loperamide. An oral dosage of 200 mg/kg body weight of a methanolic extract of *F. Nervosa* was administered to test group 1 and 400 mg/kg body weight of the same methanolic extract was administered to test group 2. Thirty minutes after the previously indicated therapy, each mouse was given 0.5 mL of castor oil orally to induce diarrhea. Every mouse was housed in a separate cage with white blotting paper covering the floor. Changing the blotting sheets every hour, the experiment was run for four hours. The total amount of diarrheal feces (watery, loose stool) that occurred during the study period was noted. After one hour, the absorbent papers were replaced with new ones. The total diarrheal feces output in the control group was used as the baseline, set at 100%.

Defecation inhibition percentage was calculated using the following formula.

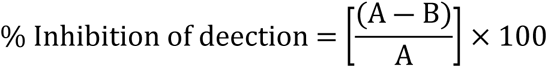

Here,

A= Mean number of defecations by castor oil;

B= Mean number of defecations by extract.

#### 2.15.2 Enteropooling assay

The effects of the plant extract on intraluminal fluid accumulation were determined using the method described by Robert *et* al. [36]. There were five groups of six mice each that were formed from the mice. Negative control groups (group 1) were treated with vehicle (distilled water for hydro methanolic extract, two milliliters each of castor oil and normal saline were given to groups 2 and 3, respectively. Groups 3 and 4 were given extracts orally at doses of 300mg/kg as well as 600 mg/kg one-to-one. An hour prior to the castor oil being administered orally animals were sacrificed by cervical dislocation ( method of cervical dislocation for euthanizing mice under 200 grams was performed without anesthesia, adhering to the 2020 AVMA and IACUC guidelines) one hour after castor oil administration, the abdomen of each animal was then opened; the small intestine was ligated at both the pyloric sphincter and the ileocecal junction and dissected, the dissected small intestine was weighed and intestinal contents were collected by carefully squeezing them into a graduated tube and volume of the contents was measured. The intestine was weighed after its contents were extracted, and the difference between the two weights recorded. Finally, the percentage reduction of intestinal secretion (volume and weight) was calculated relative to the negative control using the following formula [37].

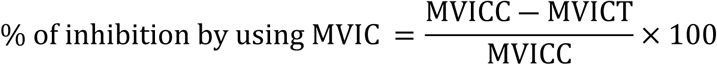

where MVIC is the mean volume of intestinal content, MVICC is the mean volume of intestinal content of the control group, and MVICT is the mean volume of intestinal content of the test group

### 2.16 Thrombolytic activity (blood clotting)

Thrombolytic assays are crucial in assessing and quantifying the efficacy of thrombolytic agents, such as tissue plasminogen activator (tPA), streptokinase, and urokinase. These drugs are widely used to treat conditions caused by blood clots, including myocardial infarction (heart attack), ischemic stroke, and pulmonary embolism [38].

#### 2.16.1 The guidelines for blood withdrawal

Guidelines for blood withdrawal from humans are established by healthcare authorities and institutions to ensure that the procedure is conducted safely, consistently, and with minimal risk to both patients and healthcare providers. Trained personnel carry out this procedure with skill and precision, following standardized protocols designed to reduce the chance of complications such as infection or injury. Key organizations, including the World Health Organization (WHO) and the Clinical and Laboratory Standards Institute (CLSI), provide comprehensive guidelines that cover every aspect of the process from patient identification and preparation to the choice of equipment and post-procedural care. These guidelines aim not only to protect the health of the volunteer’s but also to create a safe working environment for healthcare providers, emphasizing practices like proper hygiene, the use of personal protective equipment (PPE), and safe disposal of needles and other sharps [39,40]. The World University of Bangladesh’s pharmacy department’s ethics council accepted the study procedure. Before a blood sample was taken, each volunteer’s written agreement was obtained.

#### 2.16.2 Preparation of streptokinase for *in vitro* thrombolysis

Streptokinase (15, 00,000 I.U) used as a standard which was collected from Beacon pharmaceutical Ltd, Bangladesh. 5 ml sterile distilled water was added to streptokinase vial and mixed properly. From this suspension100μl (30,000 I.U) was used for *in vitro* thrombolysis [41].

#### 2.16.3 Experimental design for thrombolytic activity

The thrombolytic activity of crude extract and its different organic soluble extractives was evaluated using a technique proposed by Daginawala *et* al. with streptokinase (SK) acting as the reference material. One milliliter of distilled water was added to 10mg of methanolic extracts and their different extract components of FN, which were placed in different vials. Ten distinct pre-weighed sterile vials (1 mL/tube) holding aliquots (5 mL) of venous blood samples from healthy human volunteers who had never taken oral contraceptives were collected under aseptic circumstances. The vials were then incubated for 45 minutes at 37 °C. The weight of each vial containing clot was measured again when the clot formed and the serum was completely silent without disrupting the clot further. In each vial containing preweighed clot, 100μL aqueous solutions containing the crude extracts and different organic partitionates of FN were added separately. To the control vials, 100 μL of distilled water and 100 μL of SK were added, respectively, as negative controls and positive control. After each vial was incubated for 90 minutes at 37 °C, clot lysis was observed. Following the incubation period, the released blood serum was extracted, and vials were weighed once again to see if the clot destruction had affected the weight [42].

A percentage was calculated from the weight difference recorded prior to and following clot lysis, as shown below:

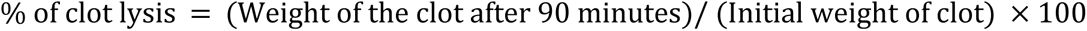

### 2.17 Statistical evaluation

The experimental data was analyzed using the Statistical Package for the Social Sciences (SPSS) version 22.0 (SPSS Inc., Chicago, IL, USA). The results were determined by taking the average of three evaluations and displaying them as mean ± standard deviation (SD). Significant deviations (p-values <0.05) between the means were determined by using the using the one-way ANOVA Duncan’s Multiple Range test (DMRT) as post-hoc.

## 3. Results

### 3.1 Analysis of phytoconstituents in the crude methanol extract of FN leaves

Our results of primary phytochemical screening identified several bioactive compounds, including steroids, tannins, saponins, phenols, alkaloids, flavonoids, and other phytocompounds in the crude methanolic extract of *F. nervosa* leaves (Table 1). In particular, we observed satisfactory amounts of phenols, flavonoids. It is well established that these bioactive compounds are beneficial for new therapeutic development of anti-inflammatory, obesity, cancer, cardiovascular diseases [43].

**Table 1.** Preliminary Phytochemical screening of crude methanolic leaves extract of *Ficus nervosa*.

| Phytochemicals | Name of the test | Remarks |
| --- | --- | --- |
| Carbohydrates | Molisch's Test | ++ |
| Reducing sugars | Fehling solution test | + |
| Tannins | Lead acetate test | +++ |
| Alkaloids | Mayer's test | ++ |
| Flavonoids | Shinoda test | +++ |
| Saponins | Frothing test | - |
| Steroids | Liebermann-Burchards test | + |
| Phenol | Ferric chloride test | +++ |

Abbreviation: Amounts shown as + = Mildly present, ++ = Moderately present, +++ = Largely present and - = Not present, depending on the depth of color.

### 3.2 Total flavonoid contents

To assess the catechin equivalent of total flavonoid levels, a complexometric technique employing aluminum chloride was utilized. Using the straight-line *Y* = 0.013 *X* + 0.1091, R2 ≥ 0.9976, which was developed from catechin (with a concentration range of 0.0 μg/mL to 160.0 μg/mL) as the standard, the flavonoid contents were evaluated in terms of catechin equivalent (Fig 1). The flavonoid concentrations of several organic extractives, including MSF, PSF, CTF, CSF, and ESF, that were extracted from FN leaves were found to be 610.20 ± 0.51, 170.01 ± 0.44, 690.30 ±61, 699.11 ± 0.10, and 639.10 ± 0.41 mg of CE/g of dry extract, respectively. The maximum number of flavonoids was observed to be highest in CSF (699.10 ± 0.10 mg of CE/g of dry extract) and lowest in PSF (170.01 ± 0.44 mg of CE/g of dry extract), followed by CSF, CTF, and MSF (Fig 2A). When compared to MSF and PSF, the flavonoid contents of ESF were considerably greater (p < 0.05)., high total phenolic content in a plant extract is a valuable indicator of its antioxidant and therapeutic potential, suggesting a broad range of health benefits and pharmaceutical applications.

**Fig 1.**
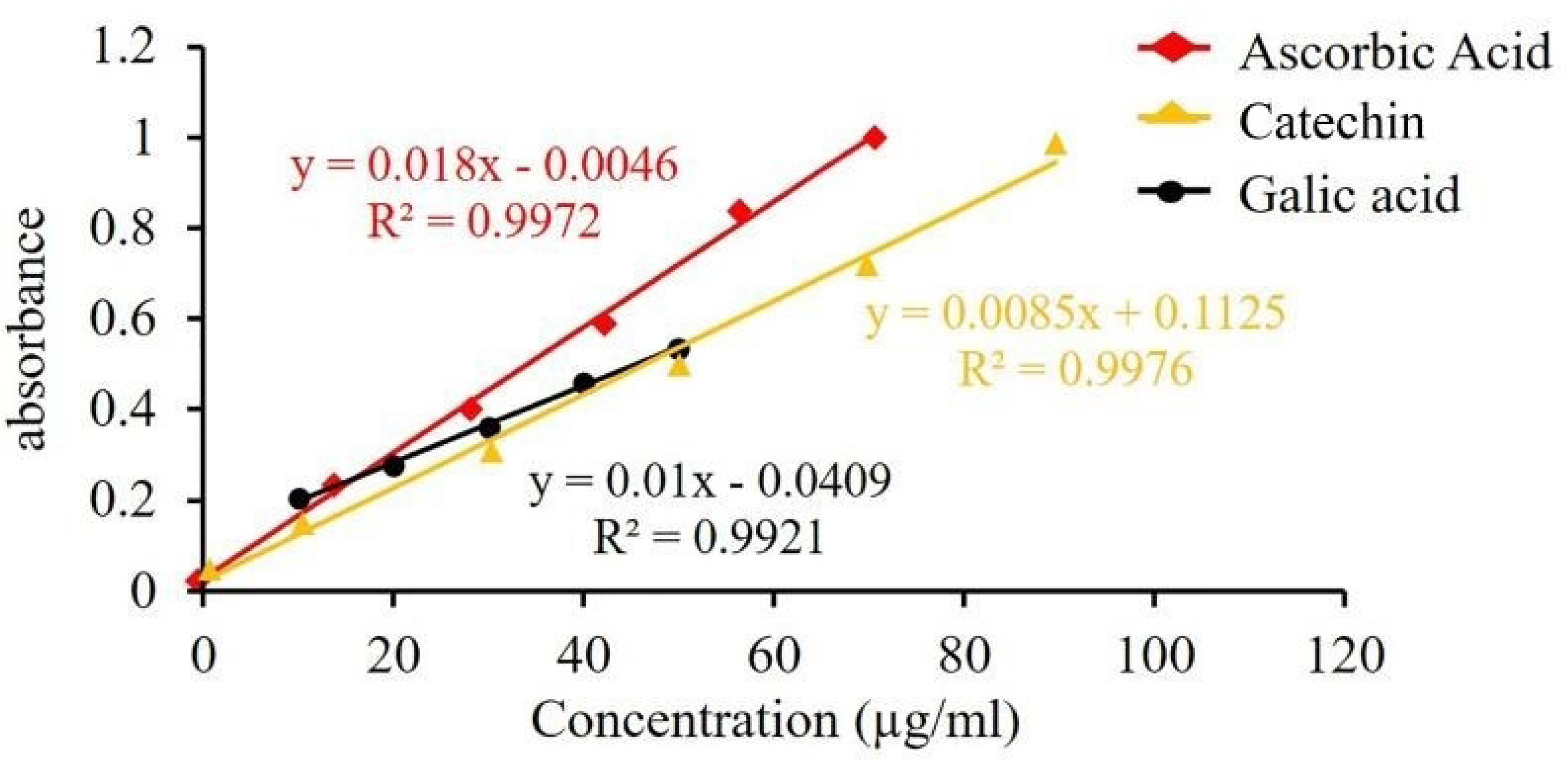
Calibration curve of ascorbic acid, catechin and gallic acid.

**Fig 2A.**
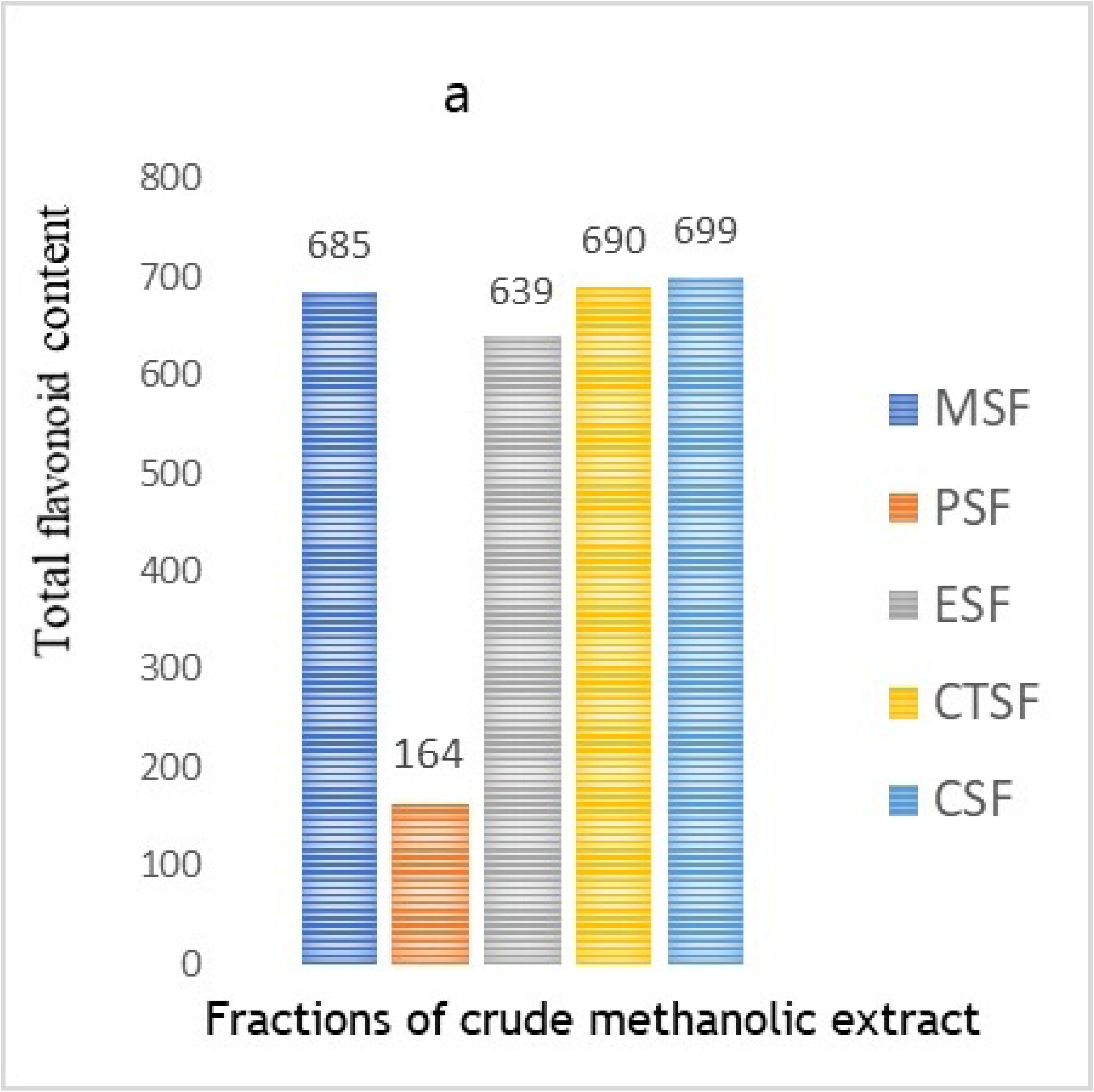
*In vitro* evaluation of fractions of leaves of *Ficus nervosa* of total flavonoid content study.

**Fig 2B.**
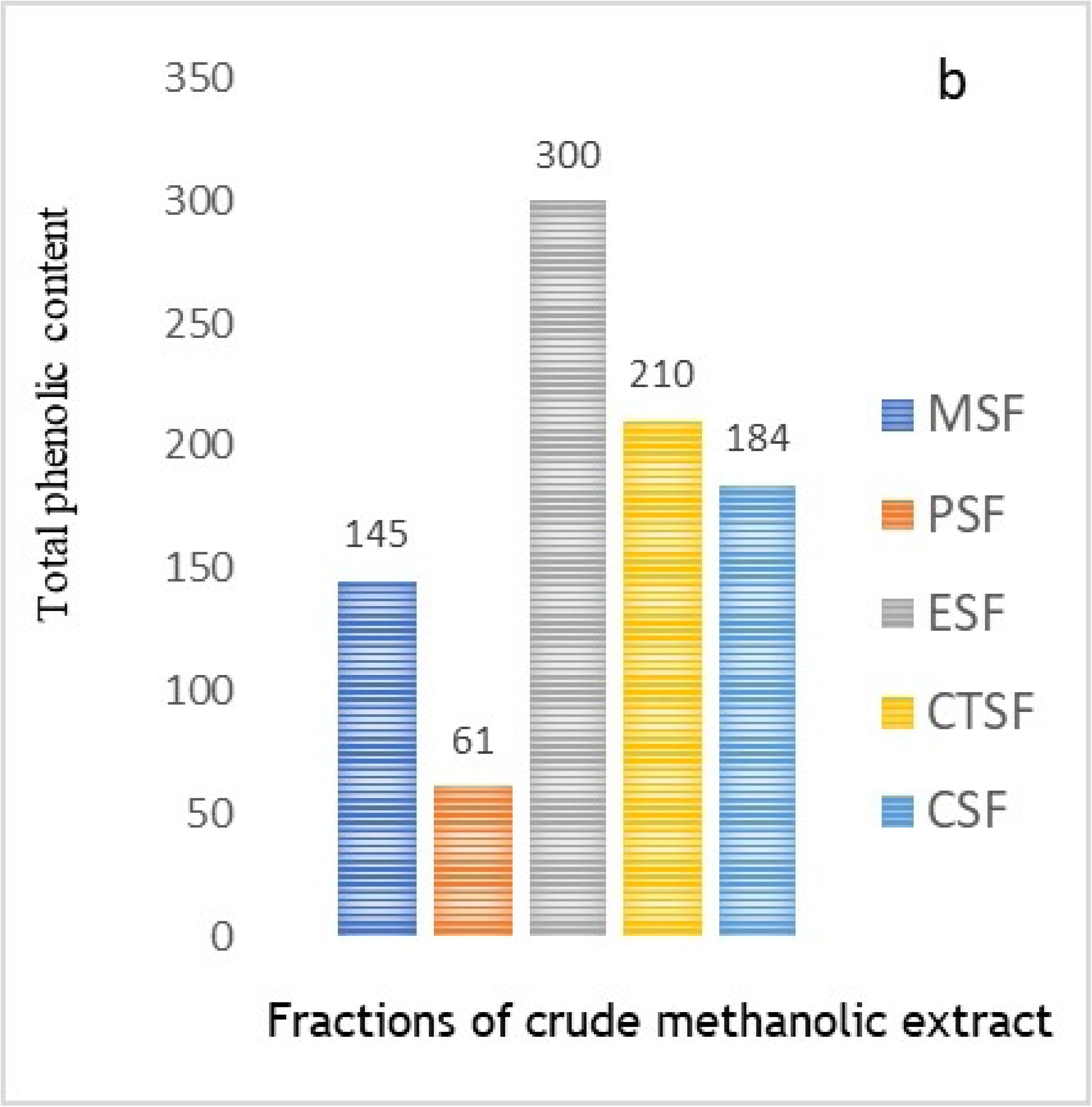
*In vitro* evaluation of fractions of leaves of *Ficus nervosa* of total phenolic content study.

### 3.3 Total Phenolic contents

The phenolic contents of the methanolic extract and its organic soluble fractionates were ascertained using the straight-line *Y* = 0.0134 *X* + 0.0461, R2 ≥ 0.9211, with gallic acid (with concentrations ranging from 0.0 μg/mL to 20.0 μg/mL) serving as the reference (Fig 1). Phenolic contents of leaves of *Ficus nervosa* extractives were found to be 148.01 ± 0.45, 69.12 ± 0.27, 220.01 ± 0.37, 194.71 ± 0.15, and 310.51 ±0.40 mg of GAE/g of MSF, PSF, CTF, CSF, and ESF, in terms of GAE/g. The results indicated that the phenolic contents were highest in ESF (310.61 ± 0.40 mg of GAE/g of dry extract) and lowest in PSF (69.11 ± 0.27 mg of GAE/g of dry extract (Fig 2B). In comparison to MSF and PSF, the phenolic contents of ESF were meaningfully higher (p < 0.05). This finding implies that all of the soluble fractions of FN leaves contain phenolic chemicals to varied degrees. The study’s findings indicate that FN leaves are a significant source of phenolic with possible health advantages.

### 3.4 Antioxidant activity

Total antioxidant capacity (TAC): In order to calculate the TAC of all organic soluble fraction in terms of ascorbic acid equivalent (AAE), a complexometric method that reduces Mo (V) to Mo (III) was used (Fig 3A). Based on a known strength of ascorbic acid, the straight line *y* = 0.18x + 0.0046, R2 = 0.9972 (Fig 1) was used to estimate the TAC in AAE. MSF, PSF, CTF, CSF, and ESF leaves of FN were found to have antioxidant capacities of 319.71 ± 0.19, 131.51 ± 0.51, 313.04 ± 0.71, 398.18 ± 0.15, and 409.90 ± 0.59 mg of AAE/g of dry extract, respectively (Table 2). Among all, CSF and ESF had the satisfactory TAC (398.18 ± 0.15, 409.91 ± 0.59 mg of AAE/g of dry extract). In both CSF and ESF, TAC was considerably (p < 0.05) higher followed by MSF, CTF and PSF.

**Fig 3A.**
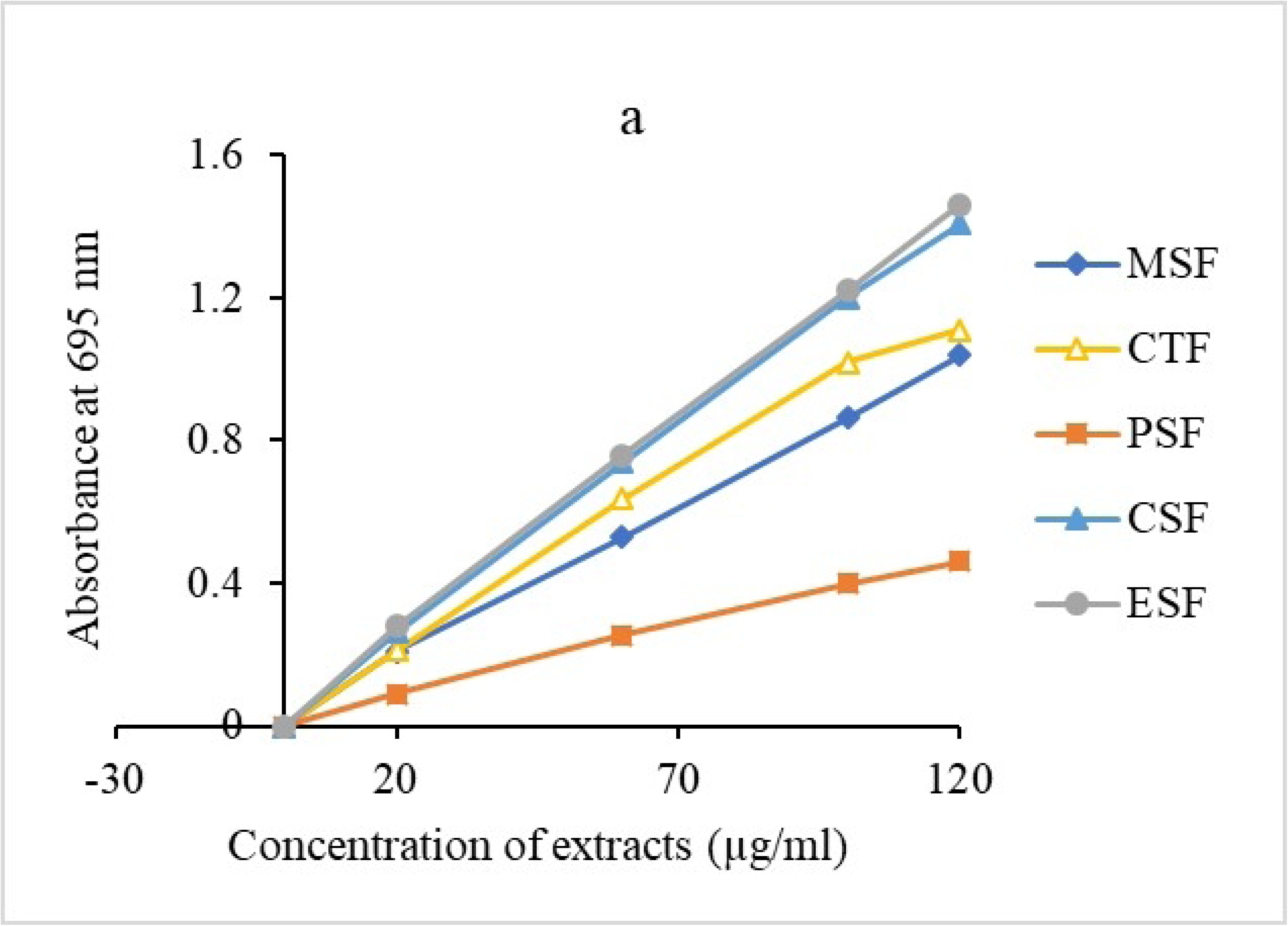
*In vitro* evaluation of antioxidant activity in different *Ficus nervosa* leaves fractions of analysis of TAC

**Fig 3B.**
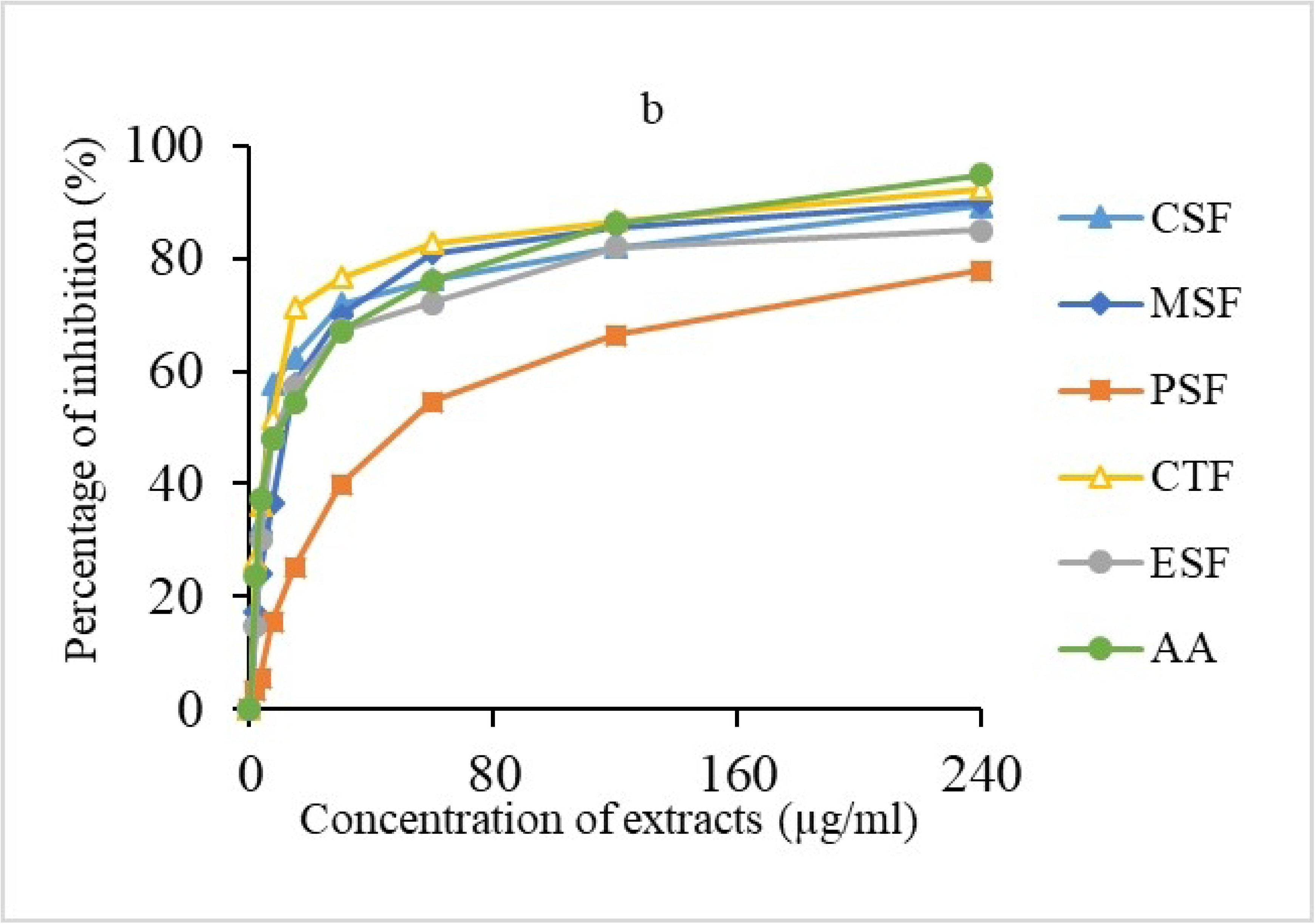
*In vitro* evaluation of antioxidant activity in different *Ficus nervosa* leaves fractions analysis of DPPH free radical scavenging activity.

**Table 2.**
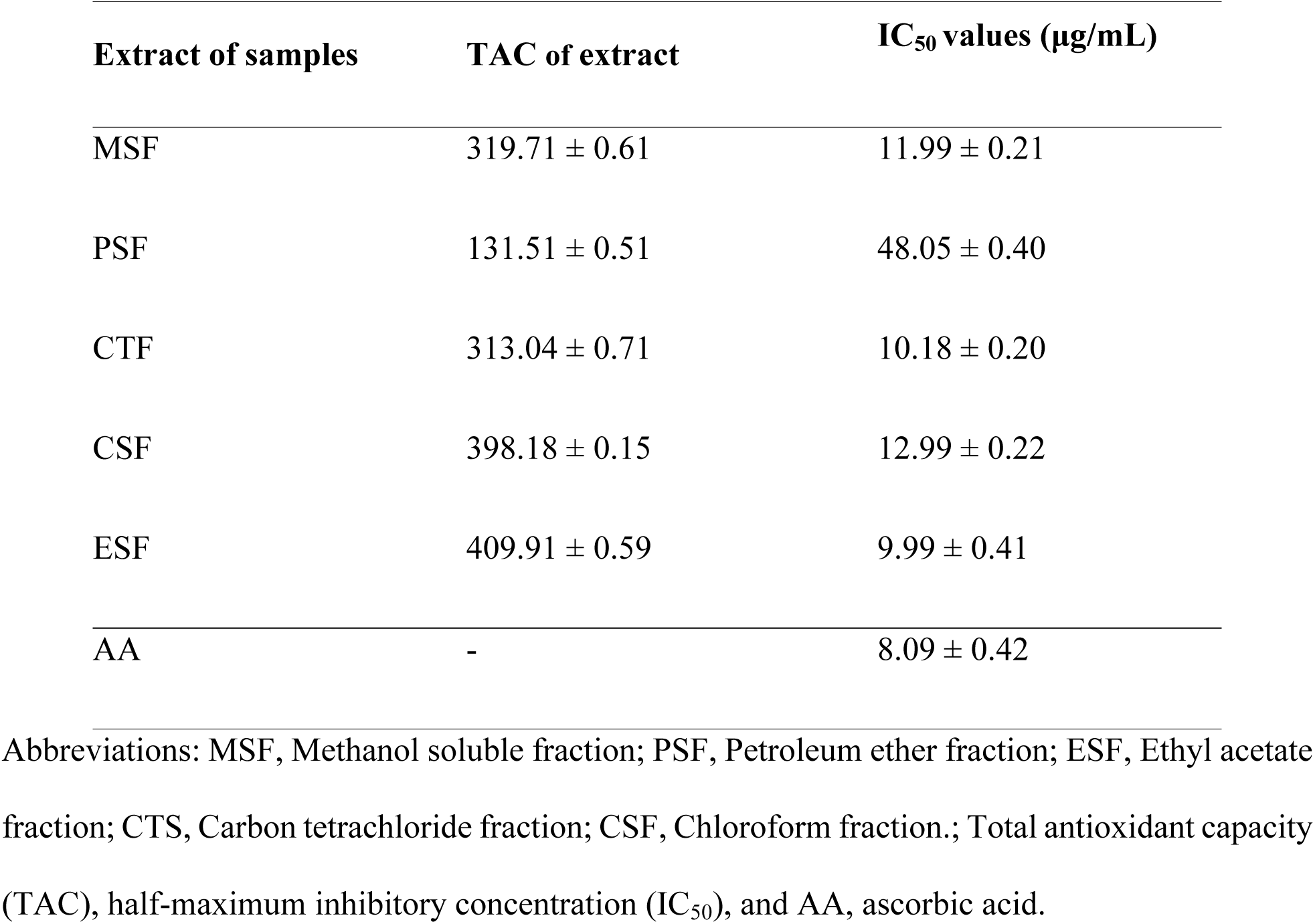
Total antioxidant capacity and DPPH free radical neutralizing property (IC_50_) of leave extractives of *Ficus nervosa*.

### 3.5 DPPH radical scavenging assay

By neutralizing the DPPH free radical, the antioxidant capacity of various extractives from FN leaves was also measured (Fig 3B). The ascorbic acid (AA), MSF, PSF, CTF, CSF, and ESF. IC_50_ values were determined using corresponding straight-line equations that were produced by graphing the % of inhibition versus the log of the sample concentration. For MSF, PSF, CTF, CSF, and ESF, the extractives of leaves of FN were found to have IC_50_ values of 11.99 ± 0.21, 48.05 ± 0.40, 10.18 ± 0.20, 12.99 ± 0.22, and 9.99 ± 0.41 μg/mL, respectively (Table 2). When compared to MSF, PSF, CSF, and AA, the IC_50_ values for ESF and CTF were meaningfully lower (p <0.05). Among the extractives ESF and CTF showed highest DPPH free radical scavenging potential (IC_50_ value 9.99 μg/mL and 10.18 μg/mL, respectively) in comparison to IC_50_ 8.09 μg/mL for ascorbic acid. From correlation analysis, a good correlation of TPC and TFC was observed with DPPH free radical scavenging (r^2^ =0.9921, 9976; p<0.05).

### 3.6 Acute toxicity

No signs of toxicity or deaths were observed after oral administration of crude methanol extract at concentration of 200, 400, 800, 1600, and 3200 mg/kg to evaluate the acute toxicity, within the first 48 hours.

In addition, no observable acute toxicity signs such as vomiting, diarrhoea, ataxia, hypoactivity, or syncope were detected. This lack of toxicity is comparable to results from earlier studies on plant-derived extracts, where comparable phytoconstituents, like phenolics and flavonoids, were found to be well-tolerated in both animal models and human applications. The absence of these adverse effects underscores the potential of the extract for further pharmacological exploration, particularly in heart diseases, algia, diarrhea and blood clotting. Previous research on similar bioactive compounds has demonstrated promising wound-healing, analgesic, diarrhoea and antimicrobial properties, further validating the extract’s potential in clinical applications [44].

### 3.7 Analgesic activity

**3.7.1 Acetic acid-induced abdominal writhes in mice**

While acetic acid was used to prepare the experiment, mice’s writhing was used to test the analgesic activity of the methanolic extract of FN leaves. In order to cause writhing and the release of endogenous molecules that excite the pain nerve and produce analgesia, acetic acid is utilized. The analgesic effects of the FN plant extract were evaluated by comparing its writhing inhibition to that of the standard drug diclofenac. The results indicated that at a dosage of 500 mg/kg, the FN extract achieved a 68.05% inhibition of acetic acid-induced writhing, while at a dosage of 250 mg/kg, it resulted in a 54.03% inhibition. In comparison, diclofenac (25 mg/kg) produced a 74.05% inhibition of the writhing response in experimental animals. Statistically, the FN extract significantly reduced the number of acetic acid-induced writhes (p<0.05) compared to the control group, although diclofenac showed a higher level of significance (p<0.01). Detailed results are presented in Table 3. Results showed that the methanolic extract of FN demonstrated a notable analgesic effect. However, more investigation is required to determine the original active component that gives FN’s methanolic extracts their analgesic effects.

**Table 3.** Statistical evaluation of *Ficus nervosa* leaves extract on acetic acid induced writhing in mice.

| Extract/ samples | No. of mice | Total writhing | Mean writhing | % of inhibition | T-test (p-values) |
| --- | --- | --- | --- | --- | --- |
| Control (Tween 80) | 5 | 160 | 31.9 | 0 | - |
| Diclofenac (25mg/kg) | 5 | 36 | 6.9 | 74.05 | 11.19 (p<.0.01) |
| Leaves Extract (FN) (250mg/kg) | 5 | 53 | 11.9 | 54.03 | 10.0 (p<.0.05) |
| Leaves Extract (FN) (500mg/kg) | 5 | 48 | 9.9 | 68.05 | 9.60 (p<.0.05) |

Abbreviation: Diclofenac as standard drug; FN is as *Ficus nervosa* leaves and
% inhibition = percent inhibition.

#### 3.7.2 Hot plate test in mice

The analgesic activity of the methanolic extract of *Ficus nervosa* leaves was evaluated using the hot plate method, with results summarized in Table 4. In the negative control group, which received only the vehicle (distilled water), no significant change in reaction time to the thermal stimulus was observed across the duration of the experiment, confirming the absence of analgesic effects in the control treatment. By contrast, administration of morphine (5 mg/kg) produced a substantial and statistically significant increase in reaction time, indicating a strong analgesic response. This effect was evident within 30 minutes post-administration, peaked around 1 hour, and while gradually diminishing over time, remained significantly elevated even at 2 hours, reflecting the sustained efficacy of morphine.

**Table 4.** Effect of methanolic leaves extract of *F. nervosa* on hot plate test in mice.

| Treatment (mg/kg) | Mean Reaction Time Response (s) ± SEM | | | | |
| --- | --- | --- | --- | --- | --- |
|  | 0min | 30min | 60min | 90min | 120min |
| DW 1 ml/kg;<br>Group 1 | 163±0.12 | 1.77±0.13 | 2.17±0.13 | 1.50±0.10 | 1.47±.12 |
| Morphine 5mg;<br>Group II | 1.66±0.22 | 5.17±0.63 | 5.15±0.27 | 6.45±0.24 | 6.47±0.23 |
| Extract FN 250 mg;<br>Group III | 1.67±0.21 | 2.17±0.21 | 3.00±0.17 | 3.00±0.20 | 2.13±0.20 |
| Extract FN 500mg;<br>Group IV | .51±0.12 | 3.17±0.17 | 3.90±0.10 | 3.90±0.37 | 3.37±0.31 |

Abbreviation: DW = Distilled Water; Morphine = standard drug; FN = *F. nervosa*, n = 6

For the methanolic extract of *Ficus nervosa*, both tested doses (250 mg/kg and 500 mg/kg) showed significant (p<0.05) increases in reaction time to the thermal stimulus at 60 and 120 minutes compared to the vehicle-treated group. These findings suggest that the extract possesses measurable analgesic properties. Notably, the analgesic effect appeared dose-dependent, with the higher dose (500 mg/kg) showing a greater prolongation of reaction time than the lower dose (250 mg/kg). However, when compared to the standard drug morphine, the analgesic effects of the extract were more pronounced in terms of both magnitude and duration. Morphine consistently produced a more potent and longer-lasting increase in pain threshold than either dose of the extract, emphasizing its superior analgesic potential in this experimental setup.

This comparison underscores the potential of *Ficus nervosa* extract as a source of analgesic compounds while highlighting the need for further research to enhance its efficacy and understand its mechanism of action relative to established analgesics like morphine.

### 3.8 Antidiarrheal test

#### 3.8.1 Caster oil induced diarrhea

As anticipated, the control group had the greatest amount of faeces (13.9 ± 2.0), whereas the conventional loperamide (3 mg/kg body weight) group had the lowest amount of faeces (2.9 ± 0.9). Defecation inhibition was seen in 71.1% of the positive control group (Table 5). During a 4-hour testing period, all of the extracts demonstrated antidiarrheal efficacy as evidenced by a decrease in the quantity of diarrheal faeces in comparison to the control group. Also encouraging was the extracts’ % of defecation inhibition. The plant leaves of FN showed 52.09% (200 mg/kg) and 59.0% (400 mg/kg) of suppression of defecation. The maximum percentage of inhibition was demonstrated by the 400 mg/kg leaves extract of FN (59.0%) between the test extracts, whereas the positive control group (loperamide at 3 mg/kg body weight) showed 73.1% suppression of defecation. Every extract showed statistically significant p values (p<0.05 and p<0.01), demonstrating its significance. One-way and Dunnett’s test were used to analyses the data.

**Table 5.** Effect of methanolic leaves extract of *Ficus nervosa* on castor oil-induced diarrhea in mice.

| Extract/Sample | Dose | Number of diarrheal faces<br>in 4 hours | % Inhibition of<br>defecation |
| --- | --- | --- | --- |
| Control | 10 mL/kg | $13.9 \pm 2.0$ | - |
| Positive<br>control(loperamide) | 3 mL/kg | $2.9 \pm 0.9$ | 73.1 |
| Leaves extract of FN<br>(Test group I) | 200 mL/kg | $6.9 \pm 0.4$ | 52.09 |
| Leaves extract of FN<br>(Test group II) | 400 mL/kg | $5.2 \pm 0.6$ | 59.0 |

Abbreviation: loperamide as positive control; FN as *Ficus nervosa*; % inhibition = percent inhibition

#### 3.8.2 Enteropooling assay

The pharmacological evaluation of enteropooling activity for the *Ficus nervosa* extract was conducted, and the results are presented in Table 6. In normal mice treated with the negative control (distilled water), the intestinal fluid volume was measured at 1.11 mL. Administration of castor oil (2 mL, orally) induced a significant (P<0.05) increase in intestinal fluid volume, reaching 2.99 mL. Pre-treatment with the FN extract an hour before castor oil administration significantly (P<0.05) attenuated the enteropooling effect, reducing intestinal fluid volume to 1.38 mL at 300 mg/kg and to 1.00 mL at 600 mg/kg. These reductions were comparable to the baseline fluid volume observed in the negative control group.

**Table 6.** Effect of *Ficus nervosa* leaves methanolic extract castor oil-induced enteropooling in mice.

| Sample/Extract | Volume of intestinal content(mL) | Weight of intestinal content (g) |
| --- | --- | --- |
| Normal saline (2ml) | 1.11 ± 0.15 | 1.01 ± 0.06 |
| Castor oil (2ml) | 2.99 ± 0.18 | 3.00 ± 0.5 |
| FN Extract (300mg/kg) + Castor oil (2ml) | 1.38 ± 0.19 | 2.00 ± 0.16 |
| FN Extract (600mg) + Castor oil (2ml) | 1.00 ± 0.18 | 1.21 ± 0.12 |

Abbreviation: FN = *Ficus nervosa*; Normal saline = negative control Castor oil = positive control

The weight of intestinal contents followed a similar trend. In the normal group, the intestinal weight was 1.01 g, while castor oil treatment substantially increased this to 3.00 g. Pre-treatment with the FN extract significantly reduced this castor oil-induced increase, with intestinal content weights decreasing from 3.00 g to 1.21 g at the highest extract dose (600 mg/kg). This reduction in both the absolute volume and weight of intestinal contents indicates a dose-dependent anti-enteropooling effect of the hydro-methanolic extract of *Ficus nervosa*.

Statistical analysis using one-way ANOVA followed by Duncan’s Multiple Range Test (DMRT) confirmed the significance of these findings (P<0.05). Data are expressed as mean ± SEM (n=4), illustrating the extract’s efficacy in mitigating castor oil-induced intestinal hypersecretion and fluid accumulation. These results highlight the potential of *Ficus nervosa* as a therapeutic agent for managing conditions involving intestinal fluid retention or hypersecretion.

### 3.9 Thrombolytic activity

Utilizing SK as a positive, which proved 69.52% blood clot lysis, the crude methanolic and other partitionates of FN were submitted to analyses the possible thrombolytic potentials. In contrast, a negative control condition of sterile distilled water investigated a small fraction of clot breaks down (3.24%). The following fractions showed the highest levels of clot lysis, represented as percentages: CSF (39.79%), MSF (39.70%), CTF (38.81%), and PSF (30.54%). Hence CSF exhibited maximum percentage of clot lysis (39.79%) in comparison of standard SK, 69.52%, Data are represented as mean ± standard deviation. In a column, means followed by the same letters (a-e) are not statistically different at a p < 0.05, as measured by the Duncan’s Multiple Range Test (DMRT), as presented in Table 7 and Fig. 4.

**Table 7.** Effect of various FN leaves extractives and streptokinase on thrombolytic activity.

| Samples | Clot lysis (%)<br>(Mean ± SEM) |
| --- | --- |
| Negative Control (Water) | 3.24 ± 0.90<sup>a</sup> |
| PSF | 30.54 ± 0.63<sup>b</sup> |
| MSF | 39.700 ± 0.44<sup>c</sup> |
| CTF | 38.81 ± 0.32<sup>d</sup> |
| CSF | 39.79 ± 0.21<sup>c</sup> |
| SK | 69.52 ± 0.11<sup>d, e</sup> |

Abbreviation: PSF, Petroleum ether fraction; MSF, methanol soluble fraction; CTS, Carbon tetrachloride fraction; CSF, Chloroform fraction. SK, Streptokinase.

**Fig 4.**
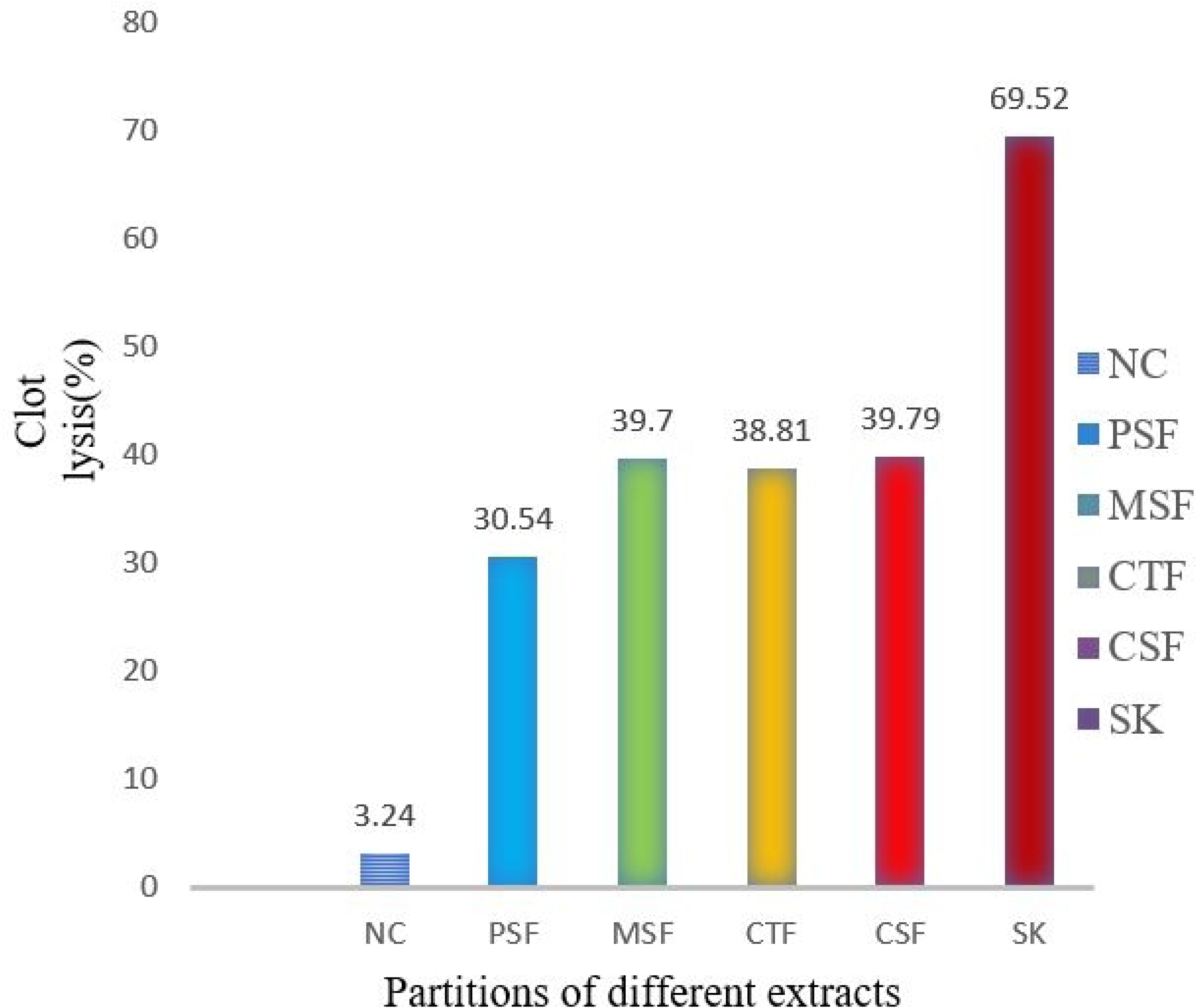
Thrombolytic properties of several *F. nervosa* leaves extractives.

## 3. Discussion

*Ficus nervosa* leaves is used in traditional medicine to treat several diseases since primitive era. To validate this folk use of the plant further we evaluated for anti-oxidant, analgesic, antidiarrheal and thrombolytic activities in our studies.

Polyphenols comprise flavonoids, phenolic acids, tannins, lignans, and coumarins, Phenols and flavonoid are naturally found phytochemical compounds in fruits, vegetables, cereals, roots, and leaves among other plant products. In this sense, recent works suggest the potential health benefits of these compounds as antioxidants against oxidative stress diseases. Special attention has been focused on phenolic compounds due to their antioxidant, antimicrobial, anti-inflammatory, anticancer, and cardiovascular protection activities [45]. In our study’s quantitative analyses ensure that naturally found phytochemical compounds contain satisfactory quantity phenol and flavonoid which encourage us to evaluate pharmacological potential of *Ficus nervosa* leaves.

The leaves extract of FN was found to good antioxidant potential. The extractives with the highest potential to scavenge free radicals caused by DPPH were CTF and ESF (IC_50_ values 10.18 ± 0.20 μg/mL and 9.99 ± 0.22, respectively), whereas ascorbic acid’s IC_50_ was 8.09 ± 0.42 μg/mL. A strong association among TPC and TFC and DPPH free radical scavenging was found through correlation analysis. Several researchers have placed their opinion that rich amounts of flavonoids and polyphenols in FN leaves also function as proton donors and have a strong association with the plant’s ability to scavenge free radical’s DPPH. The TPC and TFC are connected to the TAC of each fraction. TPC and TFC exhibited a substantial connection with TAC (r2 = 0.9921, 0.9976; p < 0.05) according to Pearson correlation analysis. Plant polyphenols and flavonoids protect the organism against oxidative stress by giving a proton to balance singlet or triplet oxygen [46].

In the acetic-acid-induced writhing test for peripheral analgesia in mice, pre-treatment 500 mg/kg and 250mg/kg b.w. of the methanolic extract and significantly reduced the writhing frequency by 68.5 % (p<.05) and 54.03% (p<.05), whereas standard diclofenac 25mg was 74.05%(p<.01). In addition to peripheral nociception, the extract of FN extract also suppressed the central nervous pain response as they significantly increased the latencies and basal pain thresholds in the hot plate tests. The acetic acid-induced abdominal writhing test is an effective screening method used to evaluate compounds for peripherally and centrally acting analgesic activities [47]. After acetic acid is administered intraperitoneally, endogenous chemicals such as PGE_2_, PGF_2α_, PGI_2_, serotonin, histamine, lipoxygenase products, and peritoneal mast cells are released and accumulate. These compounds then directly activate nociceptors, causing pain [48]. Mice’s reaction to the acetic acid is to contract their abdominal muscles, which is followed by the elongation of their body parts and the extension of their rear limbs. This constriction is believed to be mediated by local peritoneal receptors [49].

Aspirin and other NSIADs reduce acetic acid-induced writhes by delaying the synthesis and or release of these endogenous pain and inflammatory mediators [50]. Leave extract reduced the number of acetic acid-induced writhes suggesting peripherally arbitrated analgesic activity mediated by blocking the synthesis and or release of endogenous substances responsible for painful sensations. In hot plate method, mice’s reaction latency to thermally generated pain was considerably delayed by the extract. Nonetheless, morphine, the standard medication, more effectively inhibited the animals’ thermal pain response than the leaves extract. Centrally acting analgesic compounds are those that have the ability to delay the response time to a heat stimulus. [51]. Therefore, the ability of leaves extract to extend the latency period for pain suggests activity via central pain pathways, most likely by modulating endogenous substances, which are targets of pain and inflammation in addition to opioid receptors, such endogenous opioids, somatostatin, and other inhibitory hormones [52].

Castor oil-induced animal models are used to conduct *in vivo* antidiarrheal experiments. Ricinoleic acid present in caste oil which is the causes diarrhea. Ricinoleic acid changes the intestinal mucosal membrane’s electrolytic permeability by irritating and inflaming the mucosa of the small intestine. This, in turn, stimulates the intestine’s peristaltic activity. As a result, there is a decrease in the absorption of sodium and potassium ions. Endogenous prostaglandins, which excite motility and secretion, are also produced in response to these events. In addition to calculating the percentage of defecation inhibition, the total number of defecations throughout a 4-hour period was noted. In contrast to the control group, every extract exhibited antidiarrheal action in the form of decreased diarrheal faeces. Previous studies on phytochemicals found that the presence of tannins, flavonoids, alkaloids, saponins, triterpenes, and sterol in medicinal plants was linked to their antidiarrheal properties (Dash *et* al., 2014). Tanins cause the intestinal mucosa’s proteins to become denatured by producing protein tannates, which increase the mucosa’s resistance to chemical modification and reduce secretion. According to reports, flavonoids prevent the release of prostaglandins and autacoids, which may prevent castor oil-induced secretion and motility [53]. The study’s experimental leaves are abundant in tannins, alkaloids, phenols, flavonoids, steroids, and maybe additional phytochemicals. Therefore, the presence of these beneficial phytochemicals may be the cause of the plants’ strong antidiarrheal effect [54]. The leaves extracts have anti-enteropooling activity, according to the castor oil-induced enteropooling, it was statistically significant at in all parameters of extracts of FN. The anti-enteropooling activity of the extract is likely related to the presence of specific phytochemical constituents. These phytochemicals may promote the absorption of electrolytes and water, thereby mitigating the effects induced by castor oil. This mechanism may explain the observed reduction in enteropooling caused by castor oil in the leaves extract [55,56]. It was important to suggest that either reduction in intestinal motility and/or suppression of intestinal secretion was responsible for this antidiarrheal activity.

Thrombus formation one of the main vascular diseases that causes many heart conditions, particularly cardiovascular ischemic events, is thrombosis. Various thrombolytic medications, such as tissue plasminogen activator, streptokinase, and urokinase, are currently applied for thrombolysis of thrombi. However, these medications include a risk of hemorrhage and undesirable responses due to their lack of specificity. So, many efforts are ongoing over the world and trying to develop newer and specific thrombolytic agent [57]. The current investigation’s goal was to determine whether FN leaves extracts had any potential for thrombus breaking. According to our research, leaves extract’s thrombolytic activities had positive cascades when related to positive and negative controls. Hence CSF exhibited maximum percentage of clot lysis (39.79%) in comparison of standard SK, 69.52%. The results suggested that FN had phytochemicals that are in charge of the activity related to clot lysis.

## 4. Conclusion

This study highlights the significant therapeutic potential of cold methanol extracts from *Ficus nervosa* leaves, demonstrated through a comprehensive analysis of its phytochemical profile and biological activities. The presence of predominant bioactive compounds, particularly phenols and flavonoids, underpins the strong antioxidant, analgesic, antidiarrheal, and moderate thrombolytic properties observed. The extract’s potent antioxidant activity suggests that it could effectively counter oxidative stress, while its analgesic and antidiarrheal effects provide promising way for managing pain and gastrointestinal issues. Although the thrombolytic activity was moderate, it indicates potential adjunctive applications in thrombosis prevention.

These findings position *Ficus nervosa* leaves as a valuable source of bioactive compounds that may contribute to the development of natural therapeutics. Future research should focus on isolating specific active constituents and elucidating their precise mechanisms of action to facilitate the formulation of targeted, plant-based therapies. This study provides a strong foundation for further exploration, contributing valuable insights to the growing body of knowledge in ethnomedicine and modern pharmacology.

## Author Contributions

Conceptualization: Zubair Khalid Labu, Farhina Rahman Laboni Formal analysis: Zubair Khalid Labu, Samira Karim Investigation: Samira Karim, Farhina Rahman Laboni Methodology: Zubair Khalid Labu, Tarekur Rahman Project administration: Samira Karim, Md. Imran Hossain Supervision: Zubair Khalid Labu, Kaniz Fatema Writing – original draft: Zubair Khalid Labu, Sarder Arifuzzaman Writing – review & editing: Sarder Arifuzzaman, Tarekur Rahman

## Acknowledgement

The author gratefully acknowledges with sincere thanks, the Department of Pharmacy of the World University of Bangladesh provided complete laboratory facilities for this research project. Each author has approved submission and taken full responsibility for the content of the work that has been submitted.

